# Epigenetic modification and characterization of the *MGMT* promoter region using CRISPRoff in glioblastoma cells

**DOI:** 10.1101/2023.09.21.558779

**Authors:** Remi Weber, Michael Weller, Guido Reifenberger, Flavio Vasella

**Author notes:** **Corresponding author:** Flavio Vasella, MD, Departments of Neurology and Neurosurgery, Clinical Neuroscience Center, University Hospital Zurich, Rämistrasse 100, 8091 Zurich.

## Abstract

The methylation status of the O6-methylguanine DNA methyltransferase (*MGMT*) promoter region is a critical predictor of response to alkylating agents in glioblastoma. However, current approaches to study the *MGMT* status focus on analyzing models with non-identical backgrounds. Here, we present an epigenetic editing approach using CRISPRoff to introduce site-specific CpG methylation in the *MGMT* promoter region of glioma cell lines. Sanger sequencing revealed successful introduction of methylation, effectively generating differently methylated glioma cell lines with an isogenic background. The introduced methylation resulted in reduced MGMT mRNA and protein levels. Furthermore, the cell lines with *MGMT* promoter region methylation exhibited increased sensitivity to temozolomide, consistent with the impact of methylation on treatment outcomes in patients with glioblastoma. This precise epigenome-editing approach provides valuable insights into the functional relevance of *MGMT* promoter regional methylation and its potential for prognostic and predictive assessments, as well as epigenetic-targeted therapies.

## Introduction

O^6^-methylguanine DNA methyltransferase (MGMT) is a DNA repair protein, encoded by the *MGMT* gene. Its promoter region is made up of a cytosine-phosphate-guanine (CpG) island, which is a stretch of DNA containing 98 CpG sites within a region spanning approximately 1.2 kilobases around the transcription start site^1^. This CpG island has been observed to be methylated in a variety of cancers, including glioblastoma, colorectal cancer, non-small cell lung cancer, pancreatic cancer, and others^2–4^.

Aberrant methylation of the promoter region of the *MGMT* gene is of particular interest in glioblastoma patients, as it is frequently described as the most relevant predictor of response to alkylating agents such as temozolomide (TMZ)^5^. Most approaches to explore the biological role of MGMT have focused on analyzing the *MGMT* promoter region in cells from non-identical backgrounds, by means such as siRNA^6^, enzymatic inhibition^7^ as well as gene knock-out^8^.

Here we present an experimental approach based on recent advancements in epigenetic editing, namely CRISPRoff^9^, to enable the study of the impact of different *MGMT* promoter region methylation patterns in isogenic glioma cell lines.

## Results

### Targeting the *MGMT* promoter region using CRISPRoff in T-325 cells

Using a catalytically impaired CRISPR/Cas9 system conjugated to a methyltransferase, we introduced CpG site methylation specifically across the *MGMT* promoter region. This epigenetic modification was then quantified by sodium bisulfite conversion of DNA followed by Sanger sequencing of the CpG island spanning 98 CpG sites within the *MGMT* promoter region (Figure 1A). While the *MGMT* promoter region was mostly unmethylated in T-325 wildtype (WT) cells, methylation was successfully introduced across the *MGMT* promoter region, with averaged methylation levels of 36% (T-325 M.OFF) and 33% (T-325 M.OFF3), compared to 12% for T-325 WT. These results were reproduced in LN-18 cells where the averaged methylation levels were increased from 8% for LN-18 WT to 30% for the edited cells (LN-18 M.OFF). These results were corroborated for T-325 WT and T-325 M.OFF using pyrosequencing of sodium bisulfite converted DNA spanning a short segment of the *MGMT* promoter region (Figure 1B).

**Figure 1.**
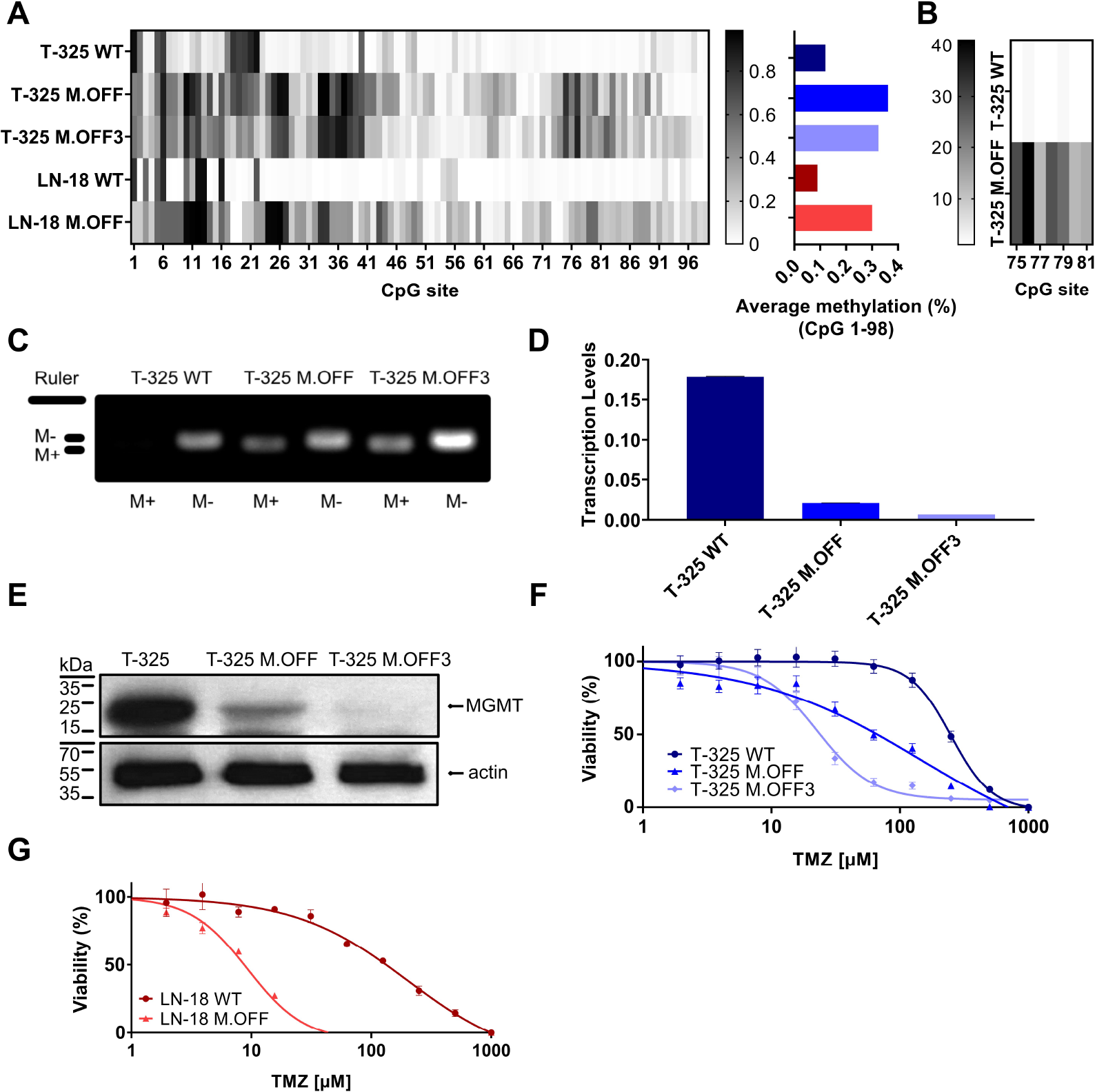
Epigenetic modification of the *MGMT* promotor region in glioma cells. A. Methylation profile of the *MGMT* promotor region, determined by Sanger sequencing of sodium bisulfite-treated DNA (EpiTect Bisulfite Kit, Qiagen) of untreated cells (T-325 WT, LN-18 WT) and cells transiently transfected with the CRISPRoff system targeting *MGMT* once (T-325 M.OFF, LN-18 M.OFF) or three times (T-325 M.OFF3) with six sgRNAs. B. Pyrosequencing of a short segment of the *MGMT* promoter region spanning 7 CpG sites in T-325 cells. C. Agarose gel of methylation-specific PCR products to quantify *MGMT* promoter methylation of the three T-325 cell lines (M+/M-: primers specific to methylated/unmethylated DNA^1^). D. Transcript levels of MGMT in T-325 cells measured by qPCR normalized to HPRT. E. MGMT protein levels in T-325 cells quantified by immunoblot. F,G. MTT assay to assess temozolomide sensitivity with and without epigenetic modification of the *MGMT* promoter region in T-325 (F) and LN-18 (G).

To further substantiate the induced change in methylation status, we analyzed the three T-325 cell lines by methylation-specific PCR (MSP), an assay commonly applied in the clinical setting^1^ (Figure 1C). Visualizing the MSP products on an agarose gel showed bands for the unmethylated PCR product in T-325 wildtype cells, while T-325 M.OFF and T-325 M.OFF3 showed bands for both unmethylated and methylated PCR products. This epigenetic modification resulted in a decrease in transcript levels in comparison to the T-325 WT cells for T-325 M.OFF, with an even more pronounced decrease in T-325 M.OFF3 (Figure 1D,E). At the same time, quantification of protein levels with immunoblot showed a reduction of the MGMT protein bands that correlates to the mRNA levels.

We next assessed the relative sensitivity to temozolomide in the T-325 and LN-18 cell lines. A strong decrease in the EC_50_ for temozolomide was observed after introducing the *MGMT* promotor region methylation for both cell lines. In T-325 cells, the TMZ EC_50_ decreased from 250 µM to 150 µM and 20 µM (T-325 WT, T-325 M.OFF, T-325 M.OFF3), while LN-18 cells experienced a similar reduction in their TMZ EC_50_ from 200 µM to 10 µM (LN-18 WT, LN-18 M.OFF) (Figure 1F,G).

### Impact of *MGMT* promoter region methylation on cell proliferation and radiosensitivity

To further elucidate the impact of the *MGMT* promoter region methylation, proliferation and radiosensitivity of modified cell lines were analyzed. The analysis of cell growth showed a significant difference between the edited and unedited cell lines with a positive correlation between degree of methylation and proliferation (Figure 2A). This was further analyzed in limited dilution conditions, where again, cells with higher methylation levels in the *MGMT* promoter region showed an increase in proliferation (Figure 2B). To determine whether the *MGMT* promoter region methylation status has an impact on radiosensitivity, the three cell lines were subjected to different doses of irradiation. While higher radiation dose led to a lower cell viability, no differences were observed between the three cell lines (Figure 2C).

**Figure 2:**
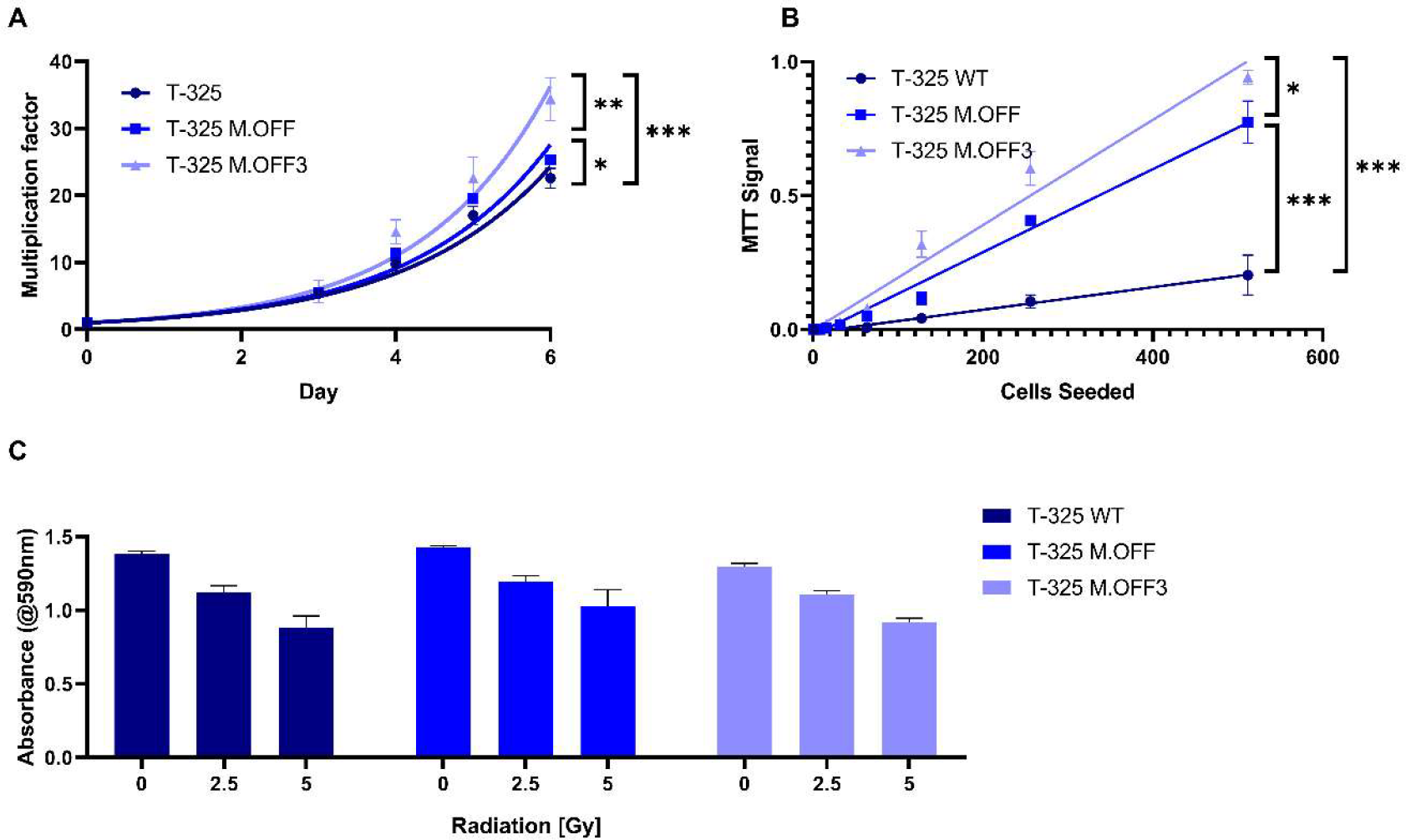
The impact of the *MGMT* promoter region methylation on cell proliferation and radiosensitivity. (A) Cell proliferation assessment of T-325 WT, M.OFF and M.OFF3 as determined by counting Hoechst 33342 stained nuclei over the course of 7 days. (B) Cell proliferation under limited dilution conditions, as determined by an MTT assay performed 10 days after seeding the specified cell number indicated on the x-axis. (C) Radiosensitivity, as determined by an MTT assay after irradiating cells with 0, 2.5 or 5 Gy twice with a 5-day interval followed by a 5-day incubation.

## Discussion

While there have been multiple attempts to overcome *MGMT*-mediated TMZ resistance in glioma cells, by means such as siRNA^6^, enzymatic inhibition^7^ or knock-out^8^, the specific induction of individual CpG methylation has so far not been recreated in a laboratory environment. Here we effectively introduced this epigenetic modification, as confirmed not just by Sanger sequencing, but also by pyrosequencing and MSP assay.

This proof of concept for the application of the CRISPRoff tool opens new opportunities for a better understanding of the methylation of the *MGMT* promoter region.

First, by the application of the epigenetic modification we demonstrated the relationship between methylation and transcript respectively protein levels in our cell lines, where both mRNA and protein levels were depleted (Fig. 1A-E). This relationship has been confirmed in cell lines^10^, but not patient tissue^11^, likely due to technical limitations resulting from the heterogeneous composition of tumor samples. Interestingly, the effects on transcript and protein levels did not correlate to the averaged methylation levels across the 98 CpG sites, potentially indicating that certain CpG sites confer a stronger impact on mRNA and protein levels than others. This result is in line with the observation, that certain regions within the *MGMT* promoter region have a stronger impact on gene expression^12^. Identifying the CpG sites that confer a stronger impact on protein levels would be especially interesting and may lead to more accurate prognostication and prediction, as current *MGMT* promoter region methylation assays only cover short segments of the *MGMT* promoter region with unclear relative impact compared to other CpG sites^13,14^.

Second, the *MGMT* methylation-dependent sensitivity to TMZ was demonstrated *in vitro* (Fig. 1 F,G). This approach enabled a direct comparison of the impact mediated solely by the methylation of the *MGMT* promoter region, circumventing added variables that are present in the literature due to intertumoral heterogeneity^12^. This observation experimentally validates the *MGMT* promoter methylation-dependent treatment outcomes observed in patients^15^, and was further substantiated in a murine glioblastoma xenograft model based on MGMT overexpression^16^.

Third, the previously discussed reduction in variables within the edited and T-325 WT cells enabled the correlation between methylation levels and phenotypic changes in proliferation as well as radiosensitivity. We showed that methylation of the *MGMT* promoter region resulted in a higher proliferation rate among the three cell lines tested (Fig. 2A,B), which has also been described in cholangiocarcinoma. In cholangiocarcinoma, *MGMT* methylation increases the number of cells entering S-phase by inhibiting p21, p27, and Cyclin E expression, which could be a possible explanation for the observed effects in the glioma cell lines^17^.

We furthermore demonstrated that the *MGMT* promoter region methylation does not correlate with radiosensitivity *in vitro* (Fig. 2C), contrary to results previously published^18^, but consistent with current concepts of the role of MGMT^19–21^.

In conclusion, we have generated insight into the impact of epigenetic modification of the *MGMT* promoter region for temozolomide sensitivity of glioma cells. In the future, this precision epigenome-editing approach could be used to determine the biological relevance of methylation of individual CpG sites, improving the prognostic and predictive value of *MGMT* promoter region methylation assessments, as well as serve as a starting point for epigenetic therapeutic interventions.

## Methods

### Cell culture and reagents

The T-325 cells were generated at the University Hospital Zurich (Zurich, Switzerland), while the LN-18 cells were kindly provided by Dr. N. de Tribolet (Lausanne, Switzerland). The cells were cultured in Dulbecco’s modified Eagle’s medium (#11965084, Gibco Life Technologies, Paisley, UK) with 2 mM L-glutamine (#25030081, Gibco Life Technologies), 100 IU/mL penicillin (#15140122, Gibco Life Technologies), 100 μg/mL streptomycin ((#15140122, Gibco Life Technologies) and 10% fetal calf serum (#16000044, Gibco Life Technologies).

### Metabolic activity assay

Culture medium was replaced with 100 µL medium containing 5 mg/mL filtered 3-(4,5-Dimethylthiazol-2-yl)-2,5-diphenyltetrazolium bromide (MTT) (#M5655, Sigma-Aldrich) and incubated for 2 h at 37°C. Subsequently, 100 µL of lysis buffer (10% SDS (#71729, Fluka, Buchs, Switzerland), 10 mM HCl (#320331, Sigma-Aldrich) in 100 mL water) were added. After overnight incubation at 37°C the absorbance was measured at a wavelength of 590 nm using a plate reader (Infinite 200 PRO, Tecan, Mannedorf, Switzerland). Blank values were obtained by measuring wells containing all components except the cells.

### Epigenetic editing

Cells were seeded at densities between 5 × 10^3^ to 20 × 10^3^ cells/well in a 96 well plate in their respective medium 24 h before transfection. Transfections were performed using TransIT™-LT1 transfection reagent (#MIR2304, Mirus Bio) using 300 ng of the CRISPR plasmid and 100 ng of each sgRNA plasmid per well of a 96 well plate. Antibiotic selection was applied two days after transfection (2 μg/mL puromycin (#ant-pr-1, Invivogen) for 3 days).

### Plasmids and sgRNAs

CRISPRoff-v2.1 was a gift from Luke Gilbert (Addgene plasmid # 167981;a Addgene, Watertown, MA, USA)^9^. The following 6 sgRNAs were used to target *MGMT*: 1. 5’-tgcgcatcctcgctggacgc-3’, 2. 5’-gacactcaccaagtcgcaaa-3’, 3. 5’-gacccggatggcccttcggc-3’, 4. 5’-acccggtcgggcgggaacac-3’, 5. 5’-gcgaggatgcgcagactgcc-3’, 6. 5’-cggctccgccccgctctaga-3’. The qPCR was run with the following primers: HPRT1 for: 5’-cattatgctgaggatttggaaagg-3’, HPRT1 rev: 5’-cttgagcacacagagggctaca-3’, MGMT for: 5’-cctggctgaatgcctatttccac-3’, MGMT rev: 5’-gcagcttccataacacctgtctg-3’.

### Evaluation of epigenetic editing

10^5^ cells were harvested for each sample. DNA was extracted and bisulfide converted using the EpiTect Bisulfide Kit (#59104, Qiagen, Venlo, Netherlands). 50 ng of the bisulfide converted, purified DNA was PCR amplified in a 20 µL reaction either using the MSP assay primers (MGMT MSP Met for: 5’-tttcgacgttcgtaggttttcgc-3’, MGMT MSP Met rev: 5’-gcactcttccgaaaacgaaacg-3’, MGMT MSP UnMet for: 5’-tttgtgttttgatgtttgtaggtttttgt-3’, MGMT MSP UnMet rev: 5’-aactccacactcttccaaaaacaaaaca-3’, annealing temperature: 49°C) or first the amplicon large (MGMT amplicon large for: 5’-ggattttaatatagtttttttggtggata-3’, MGMT amplicon large rev: 5’-gttttaggaagtttttagaagtttttg-3’, annealing temperature: 45°C) followed by the amplicon nested primers (MGMT amplicon nested for: 5’-taattttaagtagggtttggtattttgtgt-3’, MGMT amplicon nested rev: 5’-aagtgttttttaggtgttgtttagtttttt-3’, annealing temperature: 49°C) using the EpiMark® Hot Start Taq DNA Polymerase (#M0490L, New England Biolabs, Ipswich, MA, USA).

Amplicons were prepared for sequencing by ExoSAP-IT™ Express PCR Product Cleanup Reagent (#75001.4X.1.ML, Thermo Fischer Scientific) and sent to MicroSynth for Sanger sequencing using three sequencing primers (MGMT seq 1: 5’-ttaggttttggtagtgtttaggtta-3’, MGMT seq 2: 5’-gttatttggtaaattaaggtatagagtt-3’, MGMT seq 3: 5’-gcgtttttttgttttttttaggtttt-3’). The resulting .ab1 files were deconvoluted using Tracy^22^. The obtained numeric data of each base for every position were used to determine the percentage of methylation at each position.

### Proliferation assay

Cells were seeded in their respective medium in a ClearView 96 well plate (#6005182, Perkin Elmer, Waltham, MA, USA) with a quadruplicate for each timepoint. Imaging started one day after seeding. Nuclei were stained with Hoechst 33342 (0.2 μM, incubation for 2 h, #62249, Thermo Fischer Scientific), and five fields of view per well were imaged under a 4x objective in the bright and DAPI channel of a MuviCyte Live-Cell Imaging system (#HH40000000, Perkin Elmer). Cell numbers were determined by ImageJ analysis (V1.53t, National institute of health, Bethesda, MD, USA) of DAPI images.

To determine proliferation under limited dilution conditions, cells were seeded at increasingly low cell number by serial dilution with triplicates for every condition. After 10 days, viability was measured using an MTT assay.

### Immunoblot analysis

Freshly harvested cells were supplemented with phosphatase and protease inhibitor cocktail (#04906837001 and #04693132001, Sigma-Aldrich) and lysed using RIPA lysis buffer (#20-188, Merck Millipore, Burlington, VT, USA). Protein concentration in the cell lysate was quantified using the Bradford protein assay (#500-0006, Bio-Rad). 30 μg of each sample was boiled with 4x Laemmli Sample Buffer containing 10% 2-mercaptoethanol (#1610747, BioRad, Hercules, CA, USA) and run on a Mini-PROTEAN® TGX™ Precast Gel (#4561083, BioRad) and transferred to a methanol-activated nitrocellulose membrane (#10600002, Sigma-Aldrich) using a Mini Gel Tank followed by a Blot Module (#A25977 and #B1000, Thermo Fischer Scientific). Subsequently, the membrane was stained using the respective antibody (MGMT: 1:500 dilution, 4°C overnight incubation, #MA5-13506, Invitrogen, Waltham, MA, USA β-Actin: 1:5000 dilution, 20°C, 1 h incubation, #sc-47778, Santa Cruz, Dallas, TX, USA). For multiple stainings on a single membrane, Western blot Strip-It buffer (#R-03722-D50, Advansta, San Jose, CA, USA) was used. To visualize the stained proteins, SuperSignal West Femto Maximum Sensitivity Substrate (#34095, Thermo Fischer Scientific) was applied and detected using the CURIX 60 processing System (AGFA, Mortsel, Belgium).

### Methylation-specific polymerase chain reaction (MSP) assay

The PCR amplification reactions for the MSP assay (described above) were loaded onto a 3% agarose gel and ran at 80 V for 2 h. The presence of DNA bands was evaluated qualitatively using a UV lamp.

### Quantitative polymerase chain reaction (qPCR)

Total RNA was extracted from 10^6^ cells using the RNeasy Mini Kit (#74104, Qiagen) and reverse transcription was performed using the High-Capacity cDNA Reverse Transcription Kit (#4368814, Applied Biosystems, Foster City, CA, USA). The qPCR was conducted using the respective primers including normalization to HPRT1 and SYBR™ Green PCR Master Mix (#4309155, Applied Biosystems). Amplification was carried out in a QuantStudio™ 6 Flex Real-Time PCR System (#4485691, Applied Biosystems) under the following conditions: UDG activation: 50°C, 2 min; Taq polymerase activation: 95°C, 2 min; Denature 95°C, 15 sec; Anneal/Extend: 60°C, 1 min, 40 cycles.

### Radiosensitivity assay

To perform radiosensitivity assays, 5 x 10^3^ cells were placed in each well of a 96-well plate. Cells were given 24 h to attach before being irradiated (RS 2000, Radsource, Brentwood, CA, USA) with either 0, 2.5 or 5 Gy twice in 5-day intervals. After another 5 days the viability was determined by MTT assay.

### Cytotoxicity assays

A total of 5 × 10^3^ cells were placed in each well of a 96-well plate. After 24 h, they were subjected to TMZ treatment as specified for a duration of 72 h in a serum-free medium. The viability of the cells, as measured by metabolic activity, was assessed after incubation for another 3 days in serum containing medium by MTT assay.

### Statistical analysis

All data are represented as mean ± standard deviation (SD) of 3 replicates unless indicated differently. Statistical analyses were performed by multiple t-tests with the two-stage step-up method of Benjamini, Krieger and Yekutieli^23^. Significance thresholds were defined as ^*^p<0.05; ^**^p<0.01; ^***^p<0.001. Data was analyzed and visualized using Prism (GraphPad, San Diego, CA, USA).

